## Supplemental Figures 1-5 for "A big-data approach to understanding metabolic rate and response to obesity in laboratory mice"

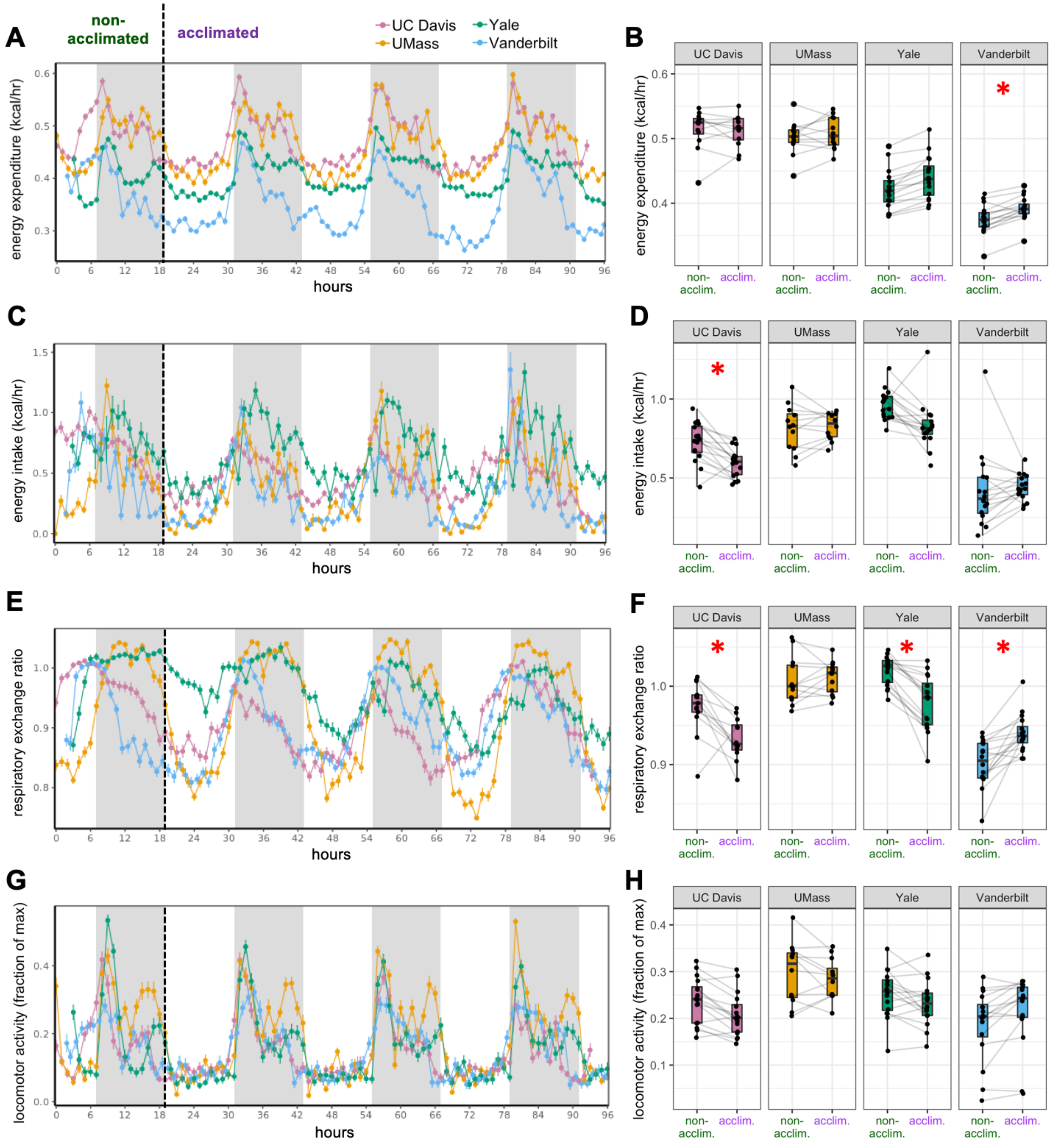


**Supplemental Figure 1. MMPC experiment: effect of acclimation.** Plots of metabolic variables vs time are shown for mice on LFD (0 weeks). EE vs time (A). Mean values for the first dark photoperiod vs the mean of the subsequent 3 dark photoperiods at each site of the MMPC experiment (B). Energy intake (C, D); RER (E, F); Locomotor activity (G, H). Values are hourly means. *, p<0.05. n = 6-8 males per group. Error bars represent SEM. Shaded regions represent the dark or active phases from 18:00 to 6:00.

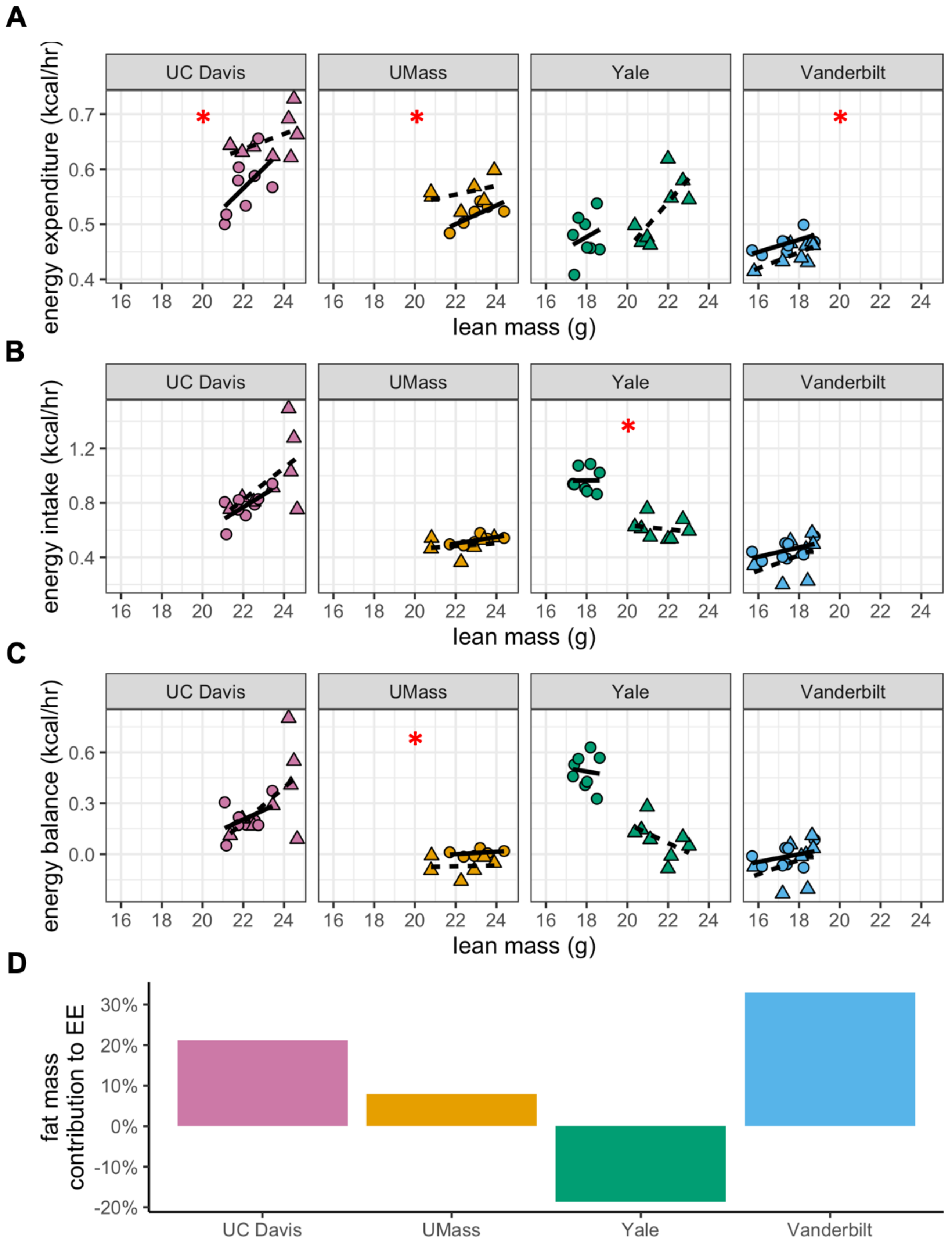


**Supplemental Figure 2. MMPC experiment: energy balance using lean mass as a covariate.** Data as in Figure 3, using the lean mass as the covariate in place of total mass. Regression plots for each of the MMPC sites vs mass. EE (A). Energy intake (B). Energy balance (C). Relative contribution of fat mass to EE for each site from a linear regression model (D). n = 6-8 males per group.

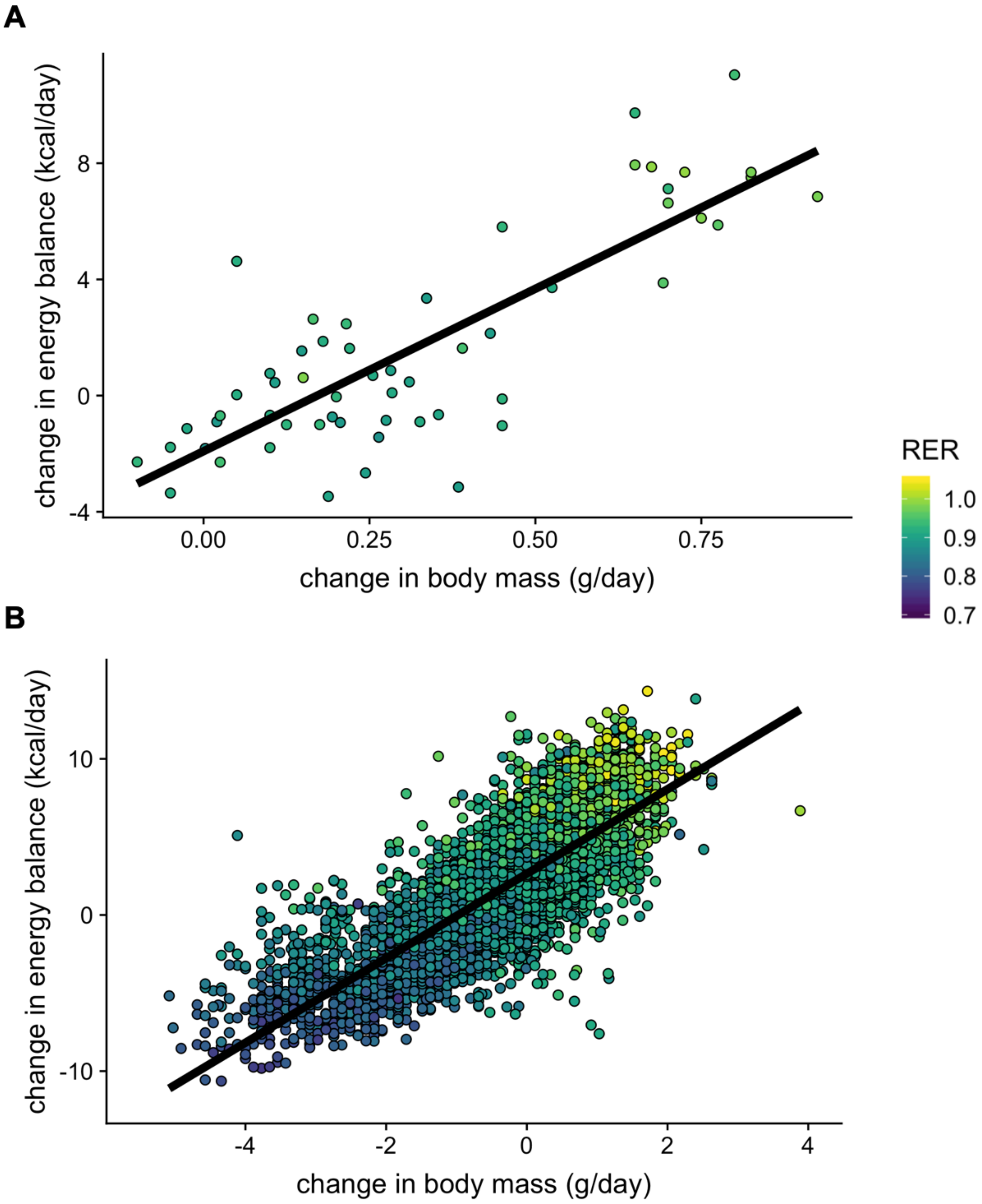


**Supplemental Figure 3. Body weight, energy balance and RER.** Plots of change in energy balance (Energy intake minus EE) vs change in body weight (mass at start of experiment minus mass at end of experiment). MMPC (n = 56 males) (A). IMPC (n = 3,781 males and 2,199 females) (B).

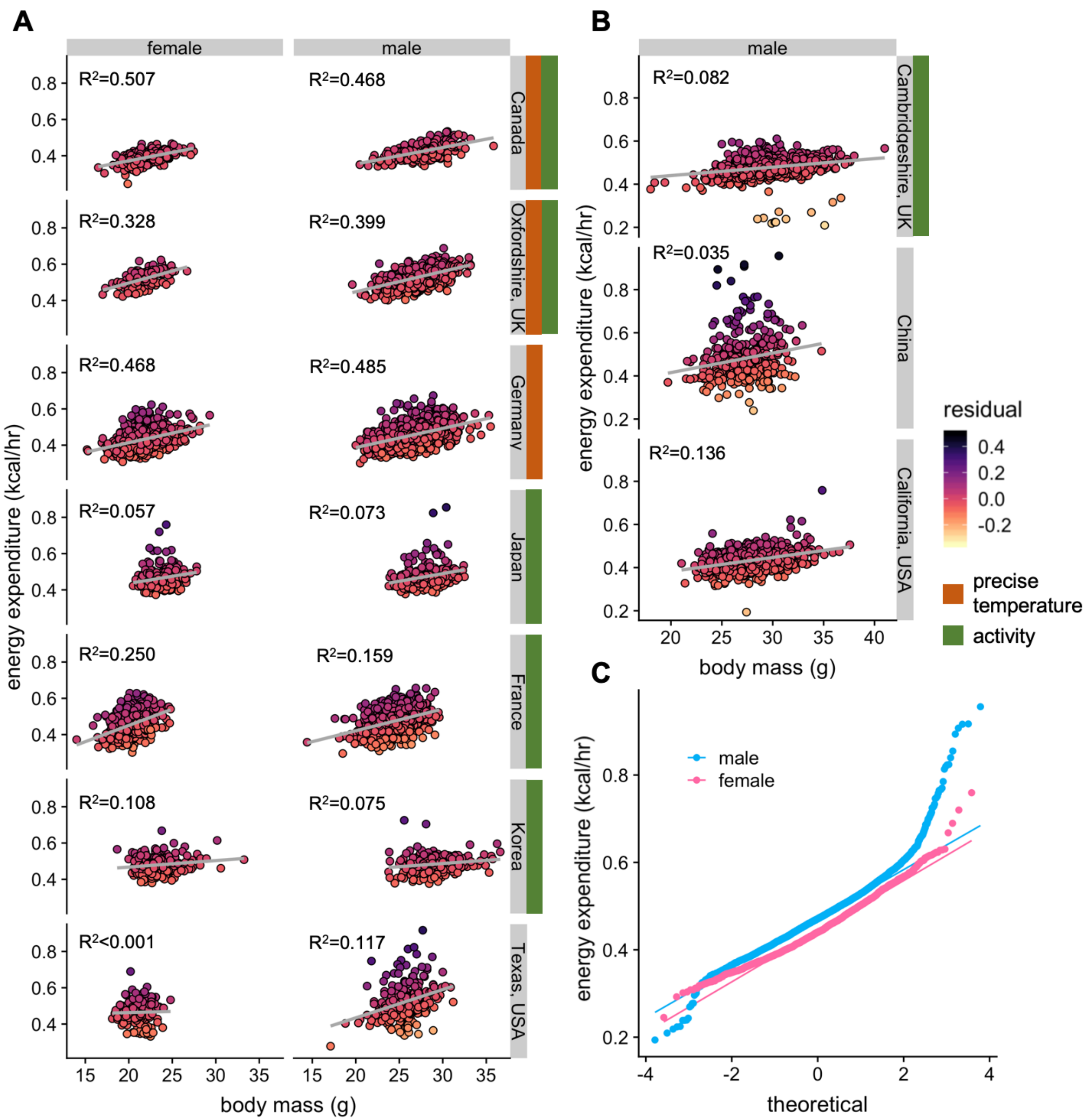


**Supplemental Figure 4. Variation in energy expenditure for WT female and male mice.** Plots of the EE vs total mass showing the quality of the fit ($R^2$ values) and shaded by residual values—the deviation from predicted values for each of the 9,545 mice (6,590 males and 2,955 females). For the 7 sites studying both sexes, the $R^2$ values were similar between female and male mice (+/- 9%). Sites with precise temperature covariate tended to have larger $R^2$ values (A). $R^2$ at the 3 sites reporting only data from male mice (B). A modified quantile-quantile plot of male and female mice from the 7 sites studying both sexes (C).

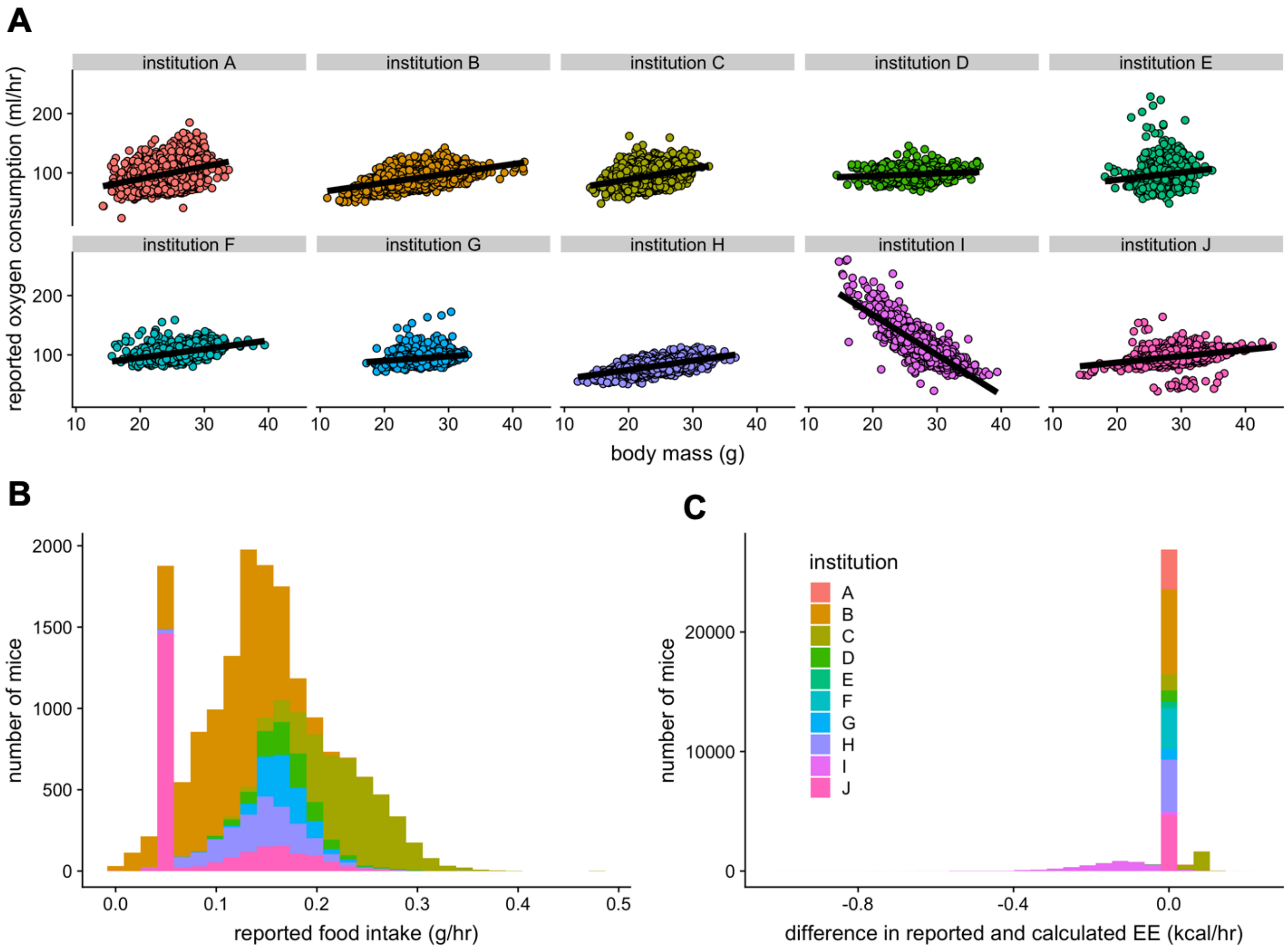


**Supplemental Figure 5. Irregularities in the IMPC dataset.** A plot of $VO_2$ vs total body mass for each of the 10 sites in the IMPC (A). Only one site demonstrates a negative mass effect due to a faulty calculation. Distribution of food intake among the reporting sites. The 0.05 g food intake was erroneously overrepresented (n = 10,021 males and 6,818 females) (B). Two sites reported EEs calculated by a different equation from the rest of the dataset (C). These 3 issues were corrected prior to analysis. n = 23,633 males and 9,646 females unless otherwise specified.
